# Pleiotropy allows recovery of phenotypic plasticity in constant environments

**DOI:** 10.1101/2020.06.02.123208

**Authors:** Enzo Kingma, Eveline T. Diepeveen, Leila Iñigo de la Cruz, Liedewij Laan

**Affiliations:** Delft University of Technology, Bionanoscience department, Van der Maasweg 9, 2629HZ Delft, the Netherlands

## Abstract

Phenotypic plasticity confers a fitness advantage to an organism by tailoring phenotype to environmental circumstances. The extent to which phenotypic plasticity emerges as an adaptive response is still unknown, however it is predicted that the emergence and maintenance of phenotypic plasticity occurs only during evolution in fluctuating environments. Interestingly, experimental studies have shown that phenotypic plasticity can be preserved for several generations during evolution in a constant environment. Here, we evolve a mutant strain of *Saccharomyces cerevisiae* that has reduced plasticity in a constant and fluctuating environment. Subsequently we compared the adaptive response of the evolved cell, both at the phenotype and genotype level. As predicted by current theory, we find that evolution in a fluctuating environment results in a recovery of phenotypic plasticity. Surprisingly, evolution in a constant environment can lead to a similar recovery of plasticity due to a pleiotropic coupling of different traits. Thus, plasticity can emerge in both fluctuating and constant environments and its prevalence may mainly be determined by network structure. In addition, pleiotropic interactions may be an important structural component of biological networks that can facilitate the recovery of phenotypic plasticity without the requirement to continuously encounter environmental fluctuations.

## Introduction

Living systems need to be able to cope with environmental changes that can negatively affect their fitness. The requirement to be able to cope with change has caused many organisms to be phenotypically plastic: they are able to adapt their characteristics in response to environmental cues, improving their probability of survival (1). Selection for phenotypic plasticity is thought to lead to the generalist phenotype, which is well adapted to a variety of different environments, but does not display the optimal phenotype for any. Conversely, a complete lack of plasticity results in the extreme specialist phenotype, which is close to the phenotypic optimum of one environment, but cannot cope with environmental changes. The observation that generalists typically do not display the specialist phenotype for any specific environment has led to conclusion that there are costs and limitations associated with plasticity (2-5).

A consequence of these costs and limitations is that the acquisition and maintenance of phenotypic plasticity is directly linked to selective pressures from the environment. A generalist only outcompetes a specialist when environmental fluctuations occur on sufficiently short timescales (6). This implies that when the environment is sufficiently stable, any generalist **c**ould potentially be replaced by a specialist that better fits the environment (7). Thus, having a plastic phenotype in a constant environment is not only non-beneficial, but is expected to be actively selected against (6). Maintaining a plastic phenotype in a constant environment therefore constitutes a significant challenge that seems to create a conflict between mutations that provide short-term fitness benefits and those that are adaptive on longer timescales.

Yet, several experimental studies have reported that the degree of phenotypic plasticity does not necessarily decrease during evolution in a constant environment (8-10). Such observations are often explained on the basis of fitness costs of plasticity, leading to the conclusion that the maintained plasticity is due to neutral or near-neutral fitness costs (9, 10). However, even if the maintenance of phenotypic plasticity in the absence of positive selection has neutral fitness costs, traits relevant for plasticity are still subject to deterioration by random mutations, as occurs during neutral drift and mutation accumulation (8, 11).

An alternative approach to understand how plasticity can be maintained without selection is to consider how plasticity relates to genetic architecture (5, 12). The topology of genetic networks is an important determinant for the accessibility of evolutionary pathways (13) and there are several examples where adaptation is restricted to only a limited set of the pathways that increase fitness (14-16). In particular, topological features can make the fitness effect of adaptive mutations in one part of the network dependent on the state of other parts of the network (17). Epistatic interactions in networks make the effect of a mutation dependent on the presence or the mutated state of other genes. Pleiotropic interactions cause a non-modular relationship between traits and the (sub)networks that regulate them. Both of these topological features have been recognized by models to play a role in the maintenance and evolution of phenotypic plasticity (12, 18).

Despite the recognized role of network topology, the genetic mechanisms that regulate plasticity and allow its maintenance in the absence of selection, are still largely unknown (2). Here, we use experimental evolution to investigate the role of environmental selection in the adaptive response of a mutant strain of *Saccharomyces cerevisiae* that has been compromised in its ability to display a plastic phenotype. Decreased phenotypic plasticity is achieved by the synthetic deletion of two genes: *BEM3* and *NRP1*. Bem3p is a known GTPase Activating Protein (GAP) of Cdc42 and is involved in polarity establishment and cytoskeletal organization (19, 20). The exact function of Nrp1p is currently unknown, but it has been implied in the formation of stress granules following glucose deprivation (21). Although deleting each gene individually has marginal effects on cell phenotype, their combined deletion has previously been found to cause a significant decrease in fitness due to defects in polarity establishment (15). We show that in addition to these polarity defects, *bem3Δnrp1Δ* cells are no longer able to properly respond to a switch from an environment with glucose as carbon source to one with ethanol as carbon source, and hence have reduced phenotypic plasticity. We expected that a variable environment, where cells repeatedly need to switch between growth on glucose and growth on ethanol, should favor phenotypes that regain some of their lost plasticity. Conversely, a constant environment in which cells only grow on glucose would have a neutral or negative effect on plasticity due to the associated costs. Surprisingly, we find that both environments readily give rise to populations that are phenotypically similar with regard to their plastic response. Whole genome-sequencing revealed that these phenotypes with similar plasticity can be traced back to mutations in two genes of the same lipid-metabolic pathway. Thus, alterations in the same pathway are found as the evolutionary solution to improved fitness regardless of the specific selective environment. We propose that through pleiotropic coupling, the effect of mutations that increase fitness in any particular environment can be extended to traits that can provide improved phenotypic plasticity.

## Results

### Pleiotropic effects of *bem3*Δ*nrp1*Δ deletion lead to decreased phenotypic plasticity

The budding yeast *S. cerevisiae* exhibits diauxic growth and preferably consumes glucose through anaerobic fermentation before switching to the aerobic respiration of ethanol (22, 23). Successful entry into the diauxic shift is hallmarked by a set of physiological changes, which include the remodeling of the actin cytoskeleton and shutdown of the Ras-cAMP-PKA pathway (23-26). As it is achieved by modifying regulatory networks in response to environmental cues, diauxic growth is a form of phenotypic plasticity. Although the deletion of *BEM3* and *NRP1* has been reported to cause defects in cell polarity, its effects on phenotypic plasticity are still uncharacterized.

We determined the degree of phenotypic plasticity of *bem3Δnrp1Δ* cells by assessing their ability to successfully pass through the diauxic shift. To this end, we quantified the population growth rates during pre- and post-diauxic growth by measuring the optical density (OD600) of a growing population over time (Figure 1A,B). We performed these growth measurements in YPD media containing a standard glucose concentration (2%, w/v) (Figure 1A) and a low glucose concentration (0.1%, w/v) (Figure 1B) for the following reason: growth in 2% glucose media provides an accurate quantification of growth in a glucose environment (Figure 1A, *T*_*Glu*_), while growth in 0.1% glucose media provides a better quantification of growth in an ethanol environment (Figure 1A, *T*_*Eth*_) (see Figure S1).

**Figure 1.**
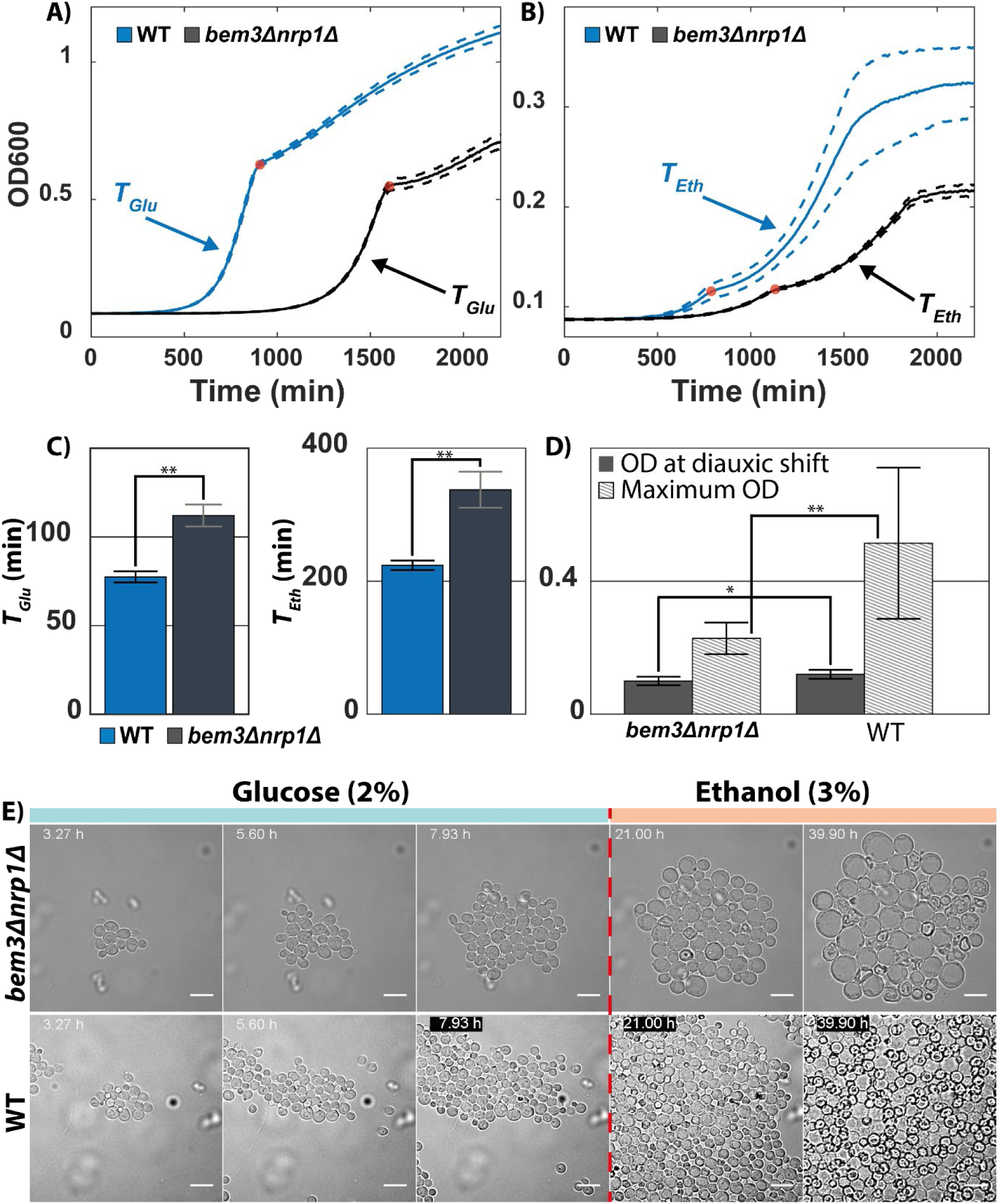
Deletion of *bem3*Δ*nrp1*Δ results in reduced phenotypic plasticity. **(A)** Growth curves of a WT (blue) and the *bem3*Δ*nrp1*Δ mutant (black) when grown in 2% glucose media. This data was used to obtain a measure for *T*_*Glu*._ Red dots indicate the point of diauxic shift, dashed lines represent the Standard Error of the Mean (SEM). **(B)** Growth curves of a WT (blue) and the *bem3*Δ*nrp1*Δ mutant (black) when grown in 0.1% glucose media. This data was used to obtain a measure for *T*_*Eth*_. Red dots indicate the point of diauxic shift, dashed lines represent the SEM. **(C)** Comparison of the values of *T*_*Glu*_ (left) and *T*_*Eth*_ (right) of the WT and the *bem3*Δ*nrp1*Δ. Error bars show the standard deviation. **(D)** Comparison of the OD at which the *bem3*Δ*nrp1*Δ mutant and the WT strain enter the diauxic shift and their OD at stationary phase when grown in YP+0.1% glucose. The plot shows that while both strains enter diauxic shift at around the same density, the final density of the populations differ. **(E)** Time-lapse series of the diauxic shift. The *bem3*Δ*nrp1*Δ and WT strain were subjected to a switch from 2% glucose media to 3% ethanol media after 8 hours in 2% glucose media (dashed red line). The images show that while the WT strain is able to resume growth, the *bem3*Δ*nrp1*Δ cells increase in size without producing daughter cells. Scale bars represent 10 µm. * = p-value<0.05, ** = p-value< 0.005, Welch t-test.

Consistent with a previous study (15), we observed that the deletion of *BEM3* and *NRP1* increases population doubling time during pre-diauxic growth on glucose (Figure 1C, Left) and found the same effect for post-diauxic growth on ethanol (Figure 1C, Right). Although slower post-diauxic growth could indicate a less efficient transition through the diauxic shift, it could also be a general effect of longer cell cycle times resulting from a defective polarization machinery. However, apart from the increase in doubling time, we also found that *bem3*Δ*nrp1*Δ cells do not reach the same density when they enter stationary phase (Figure 1B, D). We hypothesized that in addition to a defect in polarity establishment, the deletion of *bem3* and *nrp1* also causes defects in nutrient sensing. We quantified the OD values at which populations of *bem3*Δ*nrp1*Δ cells and WT cells entered diauxic shift and the subsequent stationary phase during growth in YPD containing 0.1% glucose. The difference in OD at which *bem3*Δ*nrp1*Δ cells and WT entered diauxic shift is small (∼ 1.15 fold) in comparison to the difference in final OD at stationary phase (∼2.25 fold) (Figure 1D). Thus, the ability of cells to sense the depletion of glucose prior to the diauxic shift is only mildly affected and does not appear to correlate with the lower OD at stationary phase. This indicates that a defective nutrient sensing mechanism does not fully explain the observed behavior.

We decided to investigate whether it was the response of *bem3*Δ*nrp1*Δ cells to the diauxic shift itself that leads to a decreased viability and as a consequence a lower final OD. While imaging the cells, we induced the diauxic shift under the microscope by switching from 2% glucose media to 3% ethanol media using a microfluidic device (Figure 1E). The results show a large difference in response to this induced diauxic shift: WT cells are able to continue replication after a short cessation of growth directly after the media switch (Figure 1E, bottom), while *bem3*Δ*nrp1*Δ cells are unable to resume replication in media containing 3% ethanol (Figure 1E, top). Instead, the majority of increase in biomass in the population of *bem3*Δ*nrp1*Δ cells results from a vast increase in cell size rather than the production of daughter cells. These large cells are fragile and many of them rupture as they continue to increase in size. This led us to conclude that *bem3*Δ*nrp1*Δ cells are capable of sensing and consuming nutrients when grown in 3% ethanol media, allowing them to increase in size, but unable to form daughter cells.

Other studies have shown that ethanol stress can cause depolarization of actin (27, 28). In WT budding yeast strains, this depolarization is only transient and cells are able to resume polarized growth after a short delay in the cell cycle (28). This reorganization of the actin cytoskeleton has also been observed in cells undergoing the diauxic shift after glucose exhaustion in YPD medium (24). Interestingly, failure to repolarize the actin cytoskeleton after its disassembly, for example due to severe ethanol stress, results in enlarged cells that fail to form buds (27). Due to similarities with the phenotype of the *bem3*Δ*nrp1*Δ cells, we speculate that a failure to repolarize the actin cytoskeleton underlies the inability of *bem3*Δ*nrp1*Δ cells to adapt to a switch from 2% glucose to 3% ethanol.

Based on the above observations we classify the *bem3*Δ*nrp1*Δ mutation as a pleiotropic mutation that negatively affects phenotypic plasticity. Here, we use the term pleiotropic to refer to the observation that the effect on fitness of the deletion of *BEM3* and *NRP1* depends on the environment: the fitness decrease in a 3% ethanol environment is more severe than the fitness decrease in a 2% glucose environment. Consequently, this environmental pleiotropy decreases the ability of cells to cope with a changing environment. That is, *bem3*Δ*nrp1*Δ cells are no longer able to respond to this change of carbon source in the growth media as effectively as the WT strain.

### Improved phenotypic plasticity can evolve in both a constant and a variable environment

Due to the unpredictability of natural environments, a loss of the ability to respond to environmental change will typically have large consequences for organismal fitness under natural conditions. Current theory predicts that recovery after a loss of plasticity depends on the nature of the selective environment: In the absence of environmental variation, adaptive paths that increase the mean fitness of traits under selection without recovery of plasticity are expected to be favored (2, 3, 29, 30).

Using experimental evolution, we investigated the role of the selective environment during evolutionary repair after the loss of phenotypic plasticity in *bem3*Δ*nrp1*Δ cells. We simulated a constant and changing environment by using a glucose limited continuous culture and a batch culture respectively (see Materials and Methods). In both cases only glucose was present as the initial carbon source and all ethanol was due to the fermentation of glucose during pre-diauxic growth. In a continuous culture, a constant inflow of nutrients and outflow of waste products leads to constant nutrient and cell concentrations when steady state is reached (31). Alternatively, a batch culture leads to periodic fluctuations in nutrients, waste products and cell density. As a consequence, cells in a batch culture experience a temporally heterogeneous environment, while cells in a continuous culture encounter a constant environment. In a continuous culture, cells are maintained in a state close to nutrient exhaustion, but without induction of the full stress response associated with entry into the diauxic shift (32, 33). Additionally, the constant media flow in a steady-state continuous culture results in a dilution of metabolic waste products such as ethanol. Through HPLC measurements we were able to confirm that both glucose and ethanol were present at low concentrations in our continuous culture (Table S1).

In total, we evolved 14 parallel *bem3*Δ*nrp1*Δ mutant cultures in a glucose limited continuous culture for 70 generations together with 2 WT cultures that served as controls. Based on our previous observation that the *bem3*Δ*nrp1*Δ deletion differentially affects growth in a glucose environment and an ethanol environment (Figure 1), we used the values of *T*_*Glu*_ and *T*_*Eth*_ as a first measure for fitness changes in either of these two environments. We note that because *T*_*Eth*_ specifically refers to post-diauxic growth on ethanol and requires cells to move through the diauxic shift (Figure 1B), the value of *T*_*Eth*_ is directly correlated with the ability to perform diauxic growth. Therefore, a decrease in only *T*_*Glu*_ but not *T*_*Eth*_ signifies adaptive pathways that do not positively affect plasticity, while a decrease in both *T*_*Glu*_ and *T*_*Eth*_ indicates pathways that improve plasticity.

Comparing the value of these parameters for the evolved lines to those of the ancestral strains reveals that there is a general trend in all lines evolved in a continuous culture to decrease *T*_*Glu*_ (Figure 2A), implying a constant selection for improved growth in a glucose environment. For changes in T_Eth_, two groups can be distinguished: 7 lines that adapt by increasing *T*_*Eth*_ and 7 lines that adapt by decreasing *T*_*Eth*_. The evolved *bem3*Δ*nrp1*Δ lines with the lowest (CCE1) and the highest (CCE2) value for *T*_*Eth*_ are annotated in Figure 2 and their growth curves are shown in Figure S2, togetherwith the ancestral strain. Subjecting these two evolved lines to a switch from 2% glucose media to 3% ethanol media using microfluidics (Figure 2C) confirmed that line CCE1 contains cells that show a plastic response similar to that of the WT strain. In contrast, in line CCE2 we mostly observed cells that are still unable to properly adjust to this environmental change.

**Figure 2.**
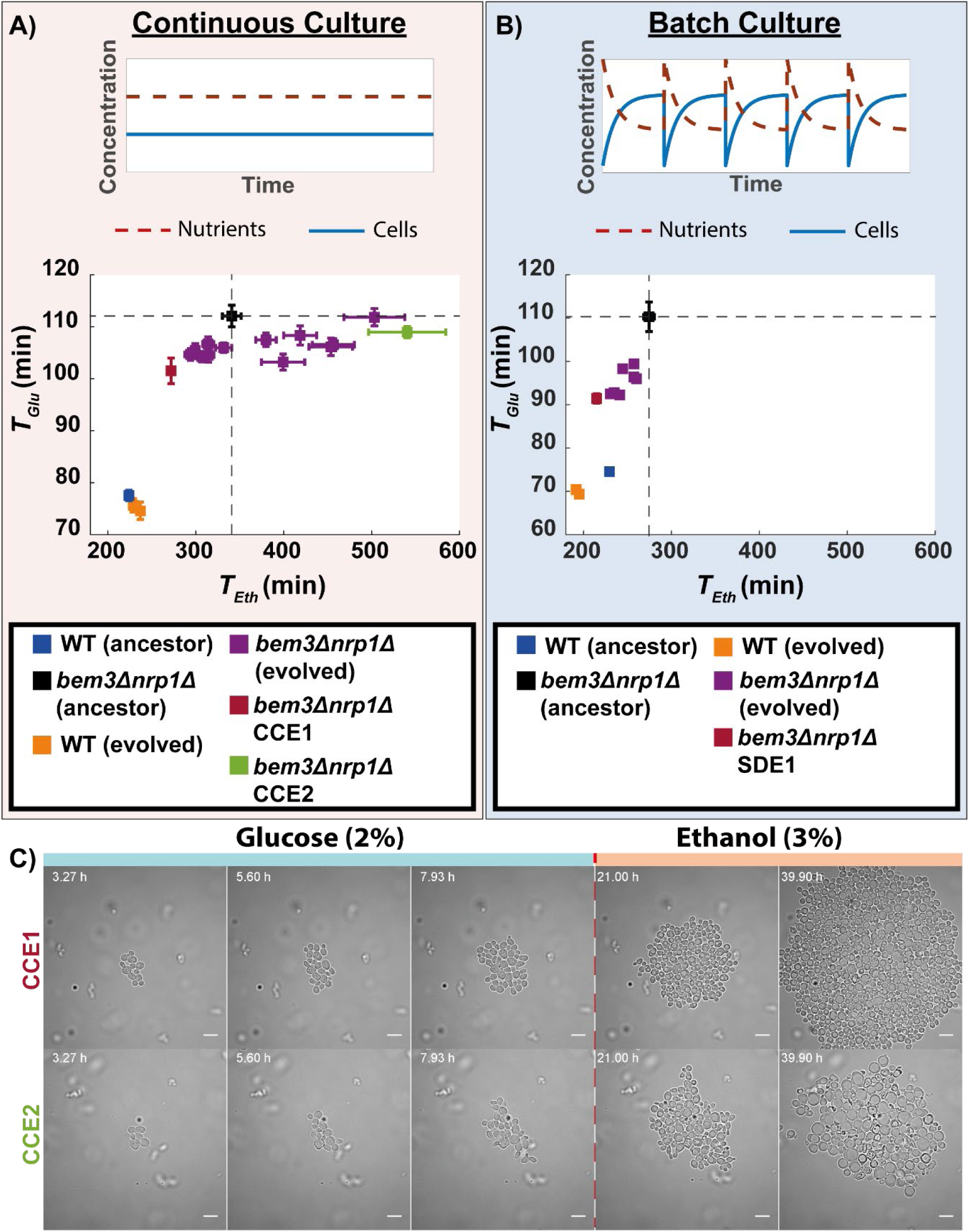
Experimental evolution of *bem3*Δ*nrp1*Δ mutants in a constant and variable environment. **(A)** (Top) In a continuous culture, both nutrient concentration and cell density remain constant over time. (Bottom) Scatter plot of *T*_*Glu*_ against *T*_*Eth*_ for 14 evolved *bem3*Δ*nrp1*Δ lines and 2 WT lines after 70 generations of evolution in a continuous culture. Dashed lines indicate the values of *T*_*Glu*_ and *T*_*Eth*_ of ancestral *bem3*Δ*nrp1*Δ strain. Error bars show the SEM. **(B)** (Top) In a batch culture there are periodic fluctuations over time in nutrient concentration and cell density. (Bottom) Scatter plot of *T*_*Glu*_ against *T*_*Eth*_ for 8 evolved *bem3*Δ*nrp1*Δ lines and 2 WT lines after 300 generations of evolution in a batch culture. Dashed lines indicate the values of *T*_*Glu*_ and *T*_*Eth*_ of ancestral *bem3*Δ*nrp1*Δ strain. Error bars show the SEM. **(C)** Time-lapse of evolved lines CCE1 and CCE2 (continuous culture) during a sudden switch from 2% glucose media to 3% ethanol media (dashed red line). The images show that evolved line CCE1 contains cells that have a response to this environmental change that is phenotypically similar to the response of the WT strain. Evolved line CCE2 has a response that resembles the response of the ancestral *bem3*Δ*nrp1*Δ, but with a smaller increase in cell size (see Figure 1E).

The evolution towards phenotypes that show reduced plasticity is in agreement with the expectation that evolution in a constant environment results in the deterioration of phenotypic plasticity, either through antagonistic pleiotropic effects or mutation accumulation (8). However, the emergence of phenotypes in our evolved *bem3*Δ*nrp1*Δ lines that adapt by decreasing both *T*_*Glu*_ and *T*_*Eth*_, indicating improved phenotypic plasticity, demonstrates that this is not a general rule. Instead, an evolutionary solution that results in increased plasticity is found in a substantial number of parallel lines (50%), which is surprising considering that random mutations with antagonistic pleiotropic effects are considered to be much more abundant than those with synergistic effects (34).

In our batch culture experiment, we evolved 8 *bem3*Δ*nrp1*Δ populations, together with 2 WT control lines. Initially, we passaged the cultures for an equal number of generations as was done for the continuous culture (70 generations). However, assessment of their fitness after 70 generations revealed that all 10 evolved populations were still phenotypically similar to their ancestors (Figure S3). After 300 generations of evolution, we exclusively observed phenotypes that decreased both parameters *T*_*Glu*_ and *T*_*Eth*_ (Figure 2B). As expected, the presence of temporal fluctuations in nutrient abundance in a batch culture environment appears to favor mutations that improve both traits and thereby phenotypic plasticity.

These results show that the two traits *T*_*Glu*_ and *T*_*Eth*_ can evolve through different couplings to each other. That is, *T*_*Eth*_ is free to both improve and deteriorate when *T*_*Glu*_ improves in environments where having a plastic phenotype does not provide a fitness benefit. Surprisingly, despite this relaxed selection on phenotypic plasticity, a substantial amount of parallel *bem3Δnrp1Δ* lines evolved in a constant environment obtain a phenotype similar to the one that evolves in a fluctuating environment. This indicates that, in our system, network architecture has a profound impact on whether the adaptive pathway will lead to enhanced phenotypic plasticity.

### Reduced phenotypic plasticity resulting from the disruption of sensing pathways is an adaptive strategy in a continuous culture

The environment determines which selection pressures are imposed on evolving populations. This difference in selection was clear in our results (Figure 2): A constant environment allows for phenotypes that reduce their plasticity, while a variable environment restricts adaptive pathways to those that improve plasticity. We wanted to determine whether the difference in selection pressure in these different environments was also visible on the molecular level. To this end, we performed Whole Genome Sequencing (WGS) on the 22 evolved *bem3*Δ*nrp1*Δ strains and the 4 evolved WT controls and compared them to the genome of their unevolved WT ancestor (see Materials and Methods). We looked for genes that were mutated in at least 2 different evolved lines to identify patterns of parallel evolution. This resulted in a total of 48 genes that acquired nonsynonymous mutations or indels.

In order to distinguish adaptations that depend on the environment from those that depend on genetic background, we grouped the mutated genes according to their occurrence (Figure 3). 37% of these mutated genes mapped to the region where all groups overlap, and thus do not hold information on adaptation to any specific selective environment or on the dependency of adaptation on genetic background. Instead, they might have been acquired through genetic hitchhiking (35) or reflect some general adaptation to long term cultivation, and we therefore dismissed these genes from further analysis. The remaining 63% of the mutated genes could be grouped in the following categories (Figure 3): (1) Environment-specific adaptation (2) Genetic background specific adaptation and (3) Adaptation to the genetic perturbation of which the effect depends on the environment.

**Figure 3.**
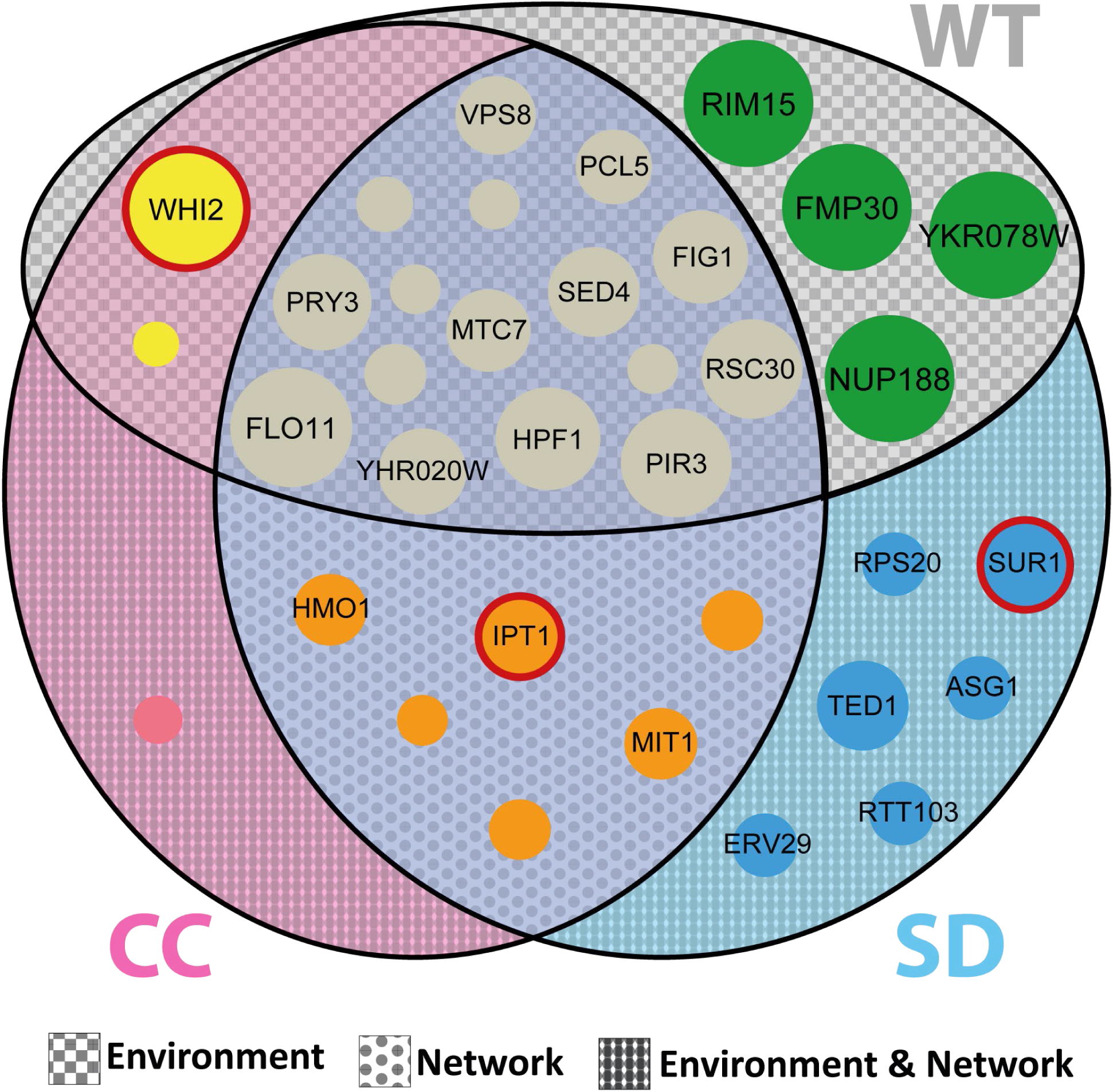
Mapping of mutations to the growth environment shows overlap in genes that are mutated in a constant and variable environment. Genes that were mutated in our *bem3Δnrp1Δ* lines during experimental evolution were mapped to a continuous culture environment (pink circle), a batch culture environment (blue circle) or overlapping regions. Mutations that were found in our control WT strains are denoted by the grey circle. The size of each node depicts the number of relevant evolved lines in which the gene was mutated. Names are shown for genes that occur in at least 25% of the lines in which it could occur.

Adaptation to the continuous culture was evident: 14/16 populations (2 WT, 12 *bem3*Δ*nrp1*Δ) acquired mutations in the stress response gene *WHI2*. The mutations occurring in *WHI2* were strongly enriched for mutations that are likely to disrupt protein function (out of 23 mutations, there was only one missense variant, all other mutations were early stop or frameshift mutations). In contrast, none of the evolved batch culture populations had acquired mutations in *WHI2* (Figure 3). This indicates that although the stress response of the diauxic shift is not fully activated during growth in a continuous culture (32, 33), pathways that sense the low concentration of the limiting nutrient may still cause growth inhibition.

In the batch culture we found no overlapping mutated genes between the WT strains and the *bem3*Δ*nrp1*Δ strain and were therefore unable to identify any genes that were obviously related to evolution in a batch culture. Surprisingly, we did find that both our WT lines had acquired the same mutation in the stress response gene *RIM15*. In particular, *RIM15* is involved in nutrient sensing (36) and has been identified as a target for inactivation during evolution in a continuous culture (37, 38). However, unlike previous studies the mutation we found in *RIM15* is not clearly disruptive to protein function. Instead, this mutation might provide some fitness benefit during growth in a variable environment, since we do find that the two WT strains evolved in a batch culture have higher fitness than their ancestor (Figure 2B).

### Populations with improved phenotypic plasticity share mutations in the same cellular pathway

We found that selection in different environments can promote the emergence of phenotypes with improved plasticity after a *bem3*Δ*nrp1*Δ deletion. We sought to explore whether the adaptations in the two environments share the same mechanism or whether they evolved distinct strategies that lead to improved phenotypic plasticity. We analyzed mutations in the genes that were mutated in the evolved *bem3*Δ*nrp1*Δ lines (evolved in a continuous or batch culture, Figure 3). These genes were grouped according to their Biological Process Gene Ontology (GO) annotation on the Saccharomyces Genome Database (Figure S4). As expected, we found that in a constant environment a larger portion of mutations occurred in genes related to the stress response than in a variable environment. Alternatively, most mutations in a variable environment occurred in genes related to transcription and translation (Figure 4). However, when a distinction is made between evolved lines that have improved phenotypic plasticity (decreased *T*_*Glu*_ and *T*_*Eth*_) and those that do not, we see that improved phenotypic plasticity correlates with an increased number of mutations in genes related to lipid metabolism. Interestingly, this trend is independent of the environment in which the populations were evolved, suggesting they might adopt the same strategy to improve plasticity.

**Figure 4.**
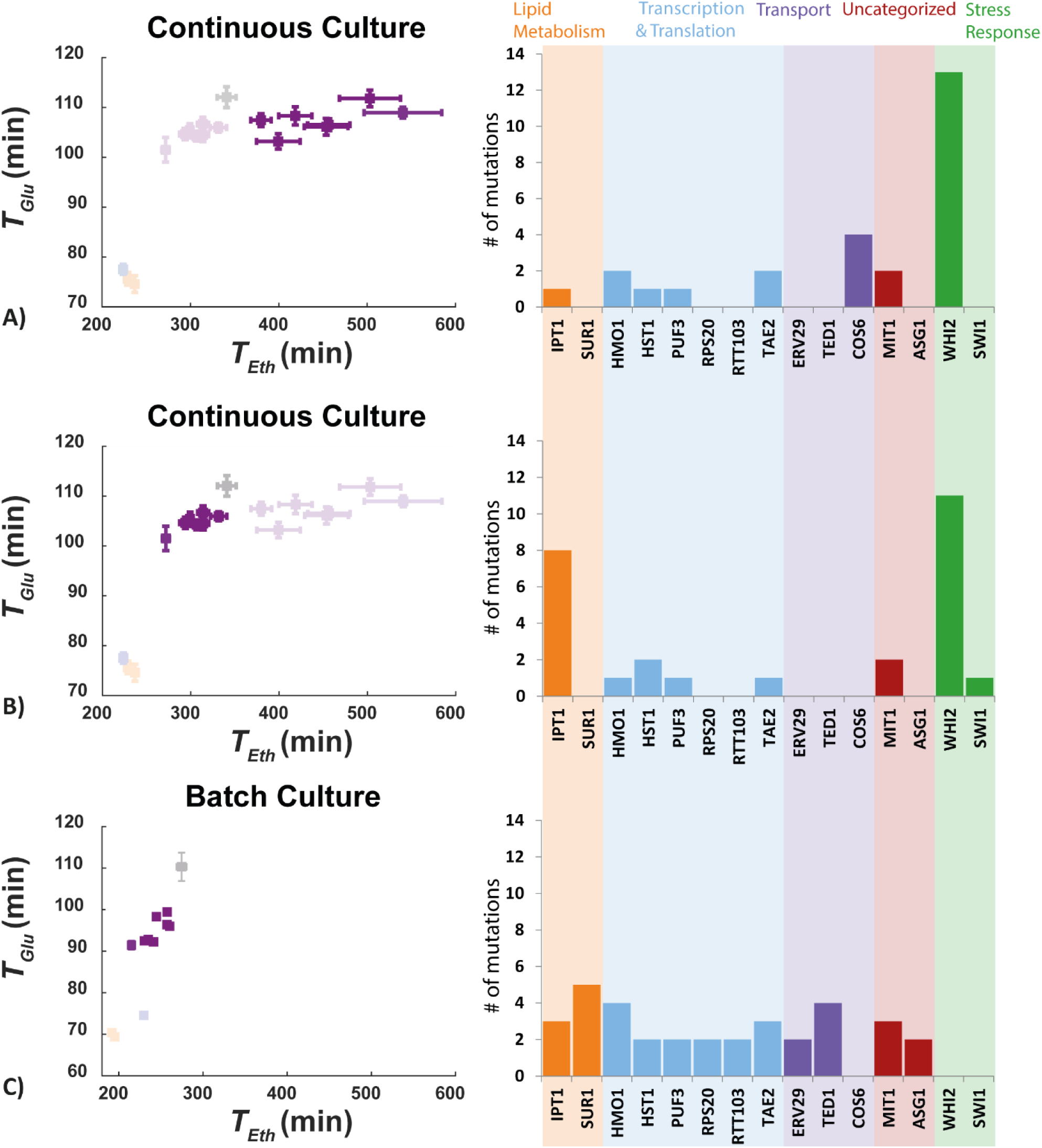
Number of mutations in each gene for different phenotypic subgroups of evolved *bem3Δnrp1Δ* lines. The number of mutant-specific mutations for (**A**) evolved continuous culture lines that decreased their respiration rate, (**B**) evolved continuous culture lines that increased their respiration rate and (**C**) evolved batch culture lines. Bars are colored according to the cellular process GO-term of the corresponding gene.

**Figure 5.**
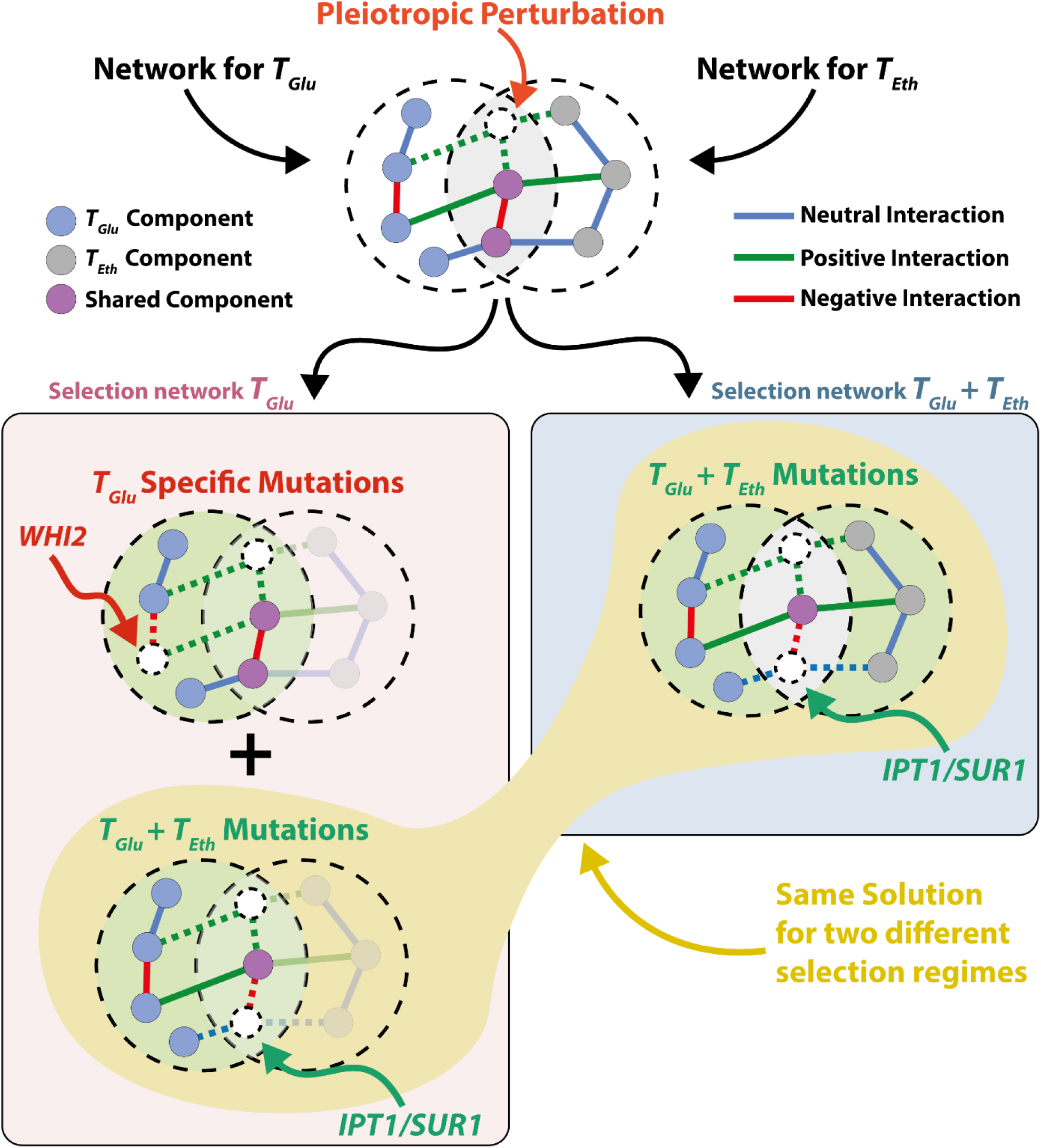
Schematic representation of the proposed mechanism through which pleiotropic interactions can promote the recovery of phenotypic plasticity. In a constant environment, only changes in the network of *T*_*Glu*_ influence fitness, while changes in the network of *T*_*Eth*_ are neutral. In a variable environment, both changes in *T*_*Glu*_ and changes in *T*_*Eth*_ affect fitness. Despite the neutral effect of *T*_*Eth*_ in a constant environment, positive pleiotropic interactions between the networks of *T*_*Glu*_ and *T*_*Eth*_ promote the fixation of mutations that improve both traits during evolution in a constant environment, resulting in recovery of phenotypic plasticity as if evolution had occurred in a variable environment.

Specifically, we found that in a continuous culture a decrease in *T*_*Eth*_ was correlated with the acquisition of mutations in *IPT1*, which encodes for an inositolphosphotransferase involved in sphingolipid synthesis (39) (Figure 4). Of the evolved *bem3*Δ*nrp1*Δ that had a decreased *T*_*Eth*_ after evolution in a constant environment, 6/7 parallel lines had acquired mutations in *IPT1* (Figure 4B), while this was the case for only 1/7 lines that had an increased *T*_*Eth*_ (Figure 4A). This suggests a causal relationship between alterations in *IPT1* and the evolution of plasticity in *bem3*Δ*nrp1*Δ mutants in a constant environment. Despite that we found a substantial number of mutations in genes related to lipid metabolism in the *bem3*Δ*nrp1*Δ populations evolved in a batch culture, only 2/8 of the parallel lines had acquired mutations in *IPT1*. Instead, the majority of the mutations in genes involved in lipid metabolism occurred in *SUR1* rather than *IPT1* (Figure 4C). Interestingly, Sur1p operates together with Csh1p directly upstream of Ipt1p in the sphingolipid synthesis pathway (39-41). In total, 5/8 parallel lines evolved in a batch culture acquired mutations in *IPT1* or *SUR1*. Thus, although adaptations in a constant and a variable environment target different genes, our data suggest that they do target the same cellular pathway for the recovery of phenotypic plasticity.

## Discussion

We experimentally tested how the ability to recover phenotypic plasticity depends on the selective nature of the environment. Our results demonstrate that phenotypic plasticity can be readily recovered after inducing a mutation that results in a loss of plasticity, regardless of whether there are temporal fluctuations in environmental conditions. We found that the phenotypes with improved plasticity in both a constant and a variable environment had acquired mutations in the same cellular pathway for complex sphingolipid synthesis. Interestingly, the ability of the sphingolipid pathway to buffer sensitivities to environmental stresses has also been described by previous studies (42, 43). These studies focus on the depolarization of the actin cytoskeleton in S*accharomyces cerevisiae* induced by salt stress. Interestingly, this appears similar to the depolarization of actin that has been found to occur during the diauxic shift (24). The deletion of either of the two amphiphysin-like proteins Rvs161p or Rvs167p from yeast makes them unable to repolarize their actin in response to salt stress (43). This phenotype is surprisingly similar to that of a *bem3*Δ*nrp1*Δ mutant during a switch from 2% glucose to 3% ethanol media, resulting in cells that can no longer form buds and increase in cell size. It was found that the deletion of either *IPT1* or *SUR1* suppresses the salt stress sensitive phenotype that results from the deletion of Rvs161p or Rvs167p (42). Thus, the same mechanism that recovers the ability of *rvs161Δ* and *rvs167Δ* mutants to cope with salts stress may also act to restore the ability of *bem3*Δ*nrp1*Δ mutants to properly respond to the diauxic shift.

Despite that several evolved populations converged at the phenotype and genotype levels, we also found mutations that appeared to be environment specific. This was most apparent in the continuous culture, where disruptive mutations in *WHI2* frequently occurred across various parallel evolved populations. Whi2p is involved in activation of the general stress response and cell cycle exit in response to nutrient depletion (26, 44). Inhibition of nutrient sensing pathways has previously been identified as the major adaptive strategy in a constant environment and mutations that inactivate Whi2p have been found in other experiments that make use of a glucose-limited continuous culture (37, 38). Disruption of nutrient sensing pathways might be beneficial in a constant environment where a response to changes in nutrient concentration is dispensable or even detrimental (38). However, such adaptations also disrupt the ability to display a plastic phenotype and decrease fitness in a variable environment. This falls in line with the conclusion that evolution in a constant environment can cause conflict between adaptations that provide a short term and a long term fitness benefit (37).

Our finding that a constant environment can promote the recovery of phenotypic plasticity contrasts with current theory. The absence of environmental fluctuations is generally expected to disfavor evolutionary pathways that lead to restored phenotypic plasticity, either due to the costs associated with plastic phenotypes or the genetic deterioration of unused traits. We show that phenotypic plasticity may be more resistant to evolutionary decay than would be expected. Our work adds to other experimental studies that have shown that phenotypic plasticity can be maintained for extended periods in non-selective environment. For example, Leiby and Marx (8) found that populations of *Escherichia coli* evolved in glucose medium improved utilization of other carbon sources after 20,000 generations and showed signs of specialization only after 50,000 generations. Similarly, Jerison, Kryazhimskiy (45) observed that adaptations in *S. cerevisiae* that increase fitness at 30°C also increase fitness at 37°C.

Several studies have linked the ability to maintain and even improve traits that are not directly under positive selection to the pleiotropic structure of the genotype to phenotype map (45-47). This contrasts with the view of pleiotropy as a source of trade-offs (48, 49) that negatively affects adaptability compared to modular network structures (50-52). Pleiotropic effects are often contextualized as a factor that impedes rather than facilitates adaptation during natural evolution. For example, Chen and Zhang (53) found that the purging of beneficial mutations due to their antagonistic pleiotropic effects on fitness in different environments decreases their fixation rate. However, trade-offs imposed by pleiotropic interactions might be most relevant for evolution when fitness is close to the pareto front (the points at which the fitness of any individual trait can only by improved by lowering the fitness of other traits) (54). Consistent with this idea, we were unable to detect signs of a trade-off with a third growth parameter in our experiments (Figure S5, S6). When the pareto front is not yet reached, synergistic interactions between traits might allow positive selection on one trait to prevent the fixation of mutations that negatively affects the other. We suggest that these pleiotropic interactions can act to delay the loss of unused traits and promote plasticity in constant environments. In other words, a pleiotropic network structure can be beneficial by extending selective pressures beyond traits that are directly under selection, at least up to the point where the pareto front is reached.

A question that remains is how mutations with synergistic effects are found so frequently amongst mutations with antagonistic effects when there is no selection for multi-trait optimization. Although there is no consensus yet (55), pleiotropic interactions are generally considered to have more antagonistic than synergistic effects (53, 56) and we do observe that mutations with antagonistic effects occur during evolution in a constant environment. In addition, for the *bem3*Δ*nrp1*Δ background it is known that adaptive pathways to higher fitness exist other than those that involve *IPT1* and *SUR1*: a previous study has shown that disruptive mutations in *BEM1* or *BEM2* are beneficial after the deletion of *BEM3* and *NRP1* (15). Despite that they would provide a significant fitness benefit, we did not find mutations in either *BEM1* or *BEM2* in any of our 22 evolved *bem3*Δ*nrp1*Δ lines. One hypothesis is that, although mutations occur randomly, they are not necessarily unbiased (57-61). In fact, mutational hotspots can affect evolutionary trajectories, as was demonstrated in a recent study that attempted to predict mutational routes in *Pseudomonas fluorescens* (62). In addition, a computational study has implicated that evolution in alternating environments can result in a network topology that causes the phenotype to be highly sensitive to mutations in specific genes (63). With regard to the evolution of phenotypic plasticity considered here, it is likely that natural strains of *S. cerevisiae* have been repeatedly selected on their ability to perform diauxic growth. We hypothesize that due to frequent positively selection on the ability to respond to environmental change, the network topology of *S. cerevisiae* might have obtained a bias for evolutionary trajectories that preserve or increase phenotypic plasticity during evolution in a constant environment. This complements another study that has reported that the evolution towards a generalist or specialist phenotype is governed by the probabilities of pleiotropic mutations being beneficial or detrimental across environments (64). These probabilities may depend on the network structures that have resulted from evolution in previously encountered environments, creating a bias.

In conclusion, we propose that a pleiotropic network structure can contribute to the maintenance of and even enhancement of phenotypic plasticity in non-fluctuating environments. Pleiotropic interactions may therefore be an important structural feature of genetic networks that prevents conflicts between mutations that provide a benefit on the short timescales with those that are beneficial on longer timescales.

## Acknowledgements

We thank Bertus Beaumont, Marjon Vos, Marco Fumasoni and Sander Tans for their critical reading and comments on the manuscript, Bas Teusink for providing feedback and useful discussions, Marc Bisschops and Nicolo Baldi for their help with the HPLC measurements and Pim America and Albert Wieland for their help with the microfluidics experiments. L.L. and E.K. gratefully acknowledges funding from the European Research Council (ERC) under the European Union’s Horizon 2020 research and innovation programme (Grant agreement No. 758132). L.L., E.T.D and LIC acknowledge support for their work from the Netherlands Organization for Scientific Research (Nederlandse Organisatie voor Wetenschappelijk Onderzoek; NWO) through a VIDI grant (016.Vidi.171.060).

## Materials and Methods

### Yeast Strains and Media Preparation

All strains used in this study are derived from the W303 background and are *MATa* haploid cells. We used yLL132 as our WT strain and yLL143a as our *bem3*Δ*nrp1*Δ strain (Laan et al., 2015), which has the same genetic background as yLL132a, but with *BEM3* and *NRP1* replaced with respectively the natMX4 (clonNAT-Nourseothricin resistance) and hphMX4 (Hygromicin B resistance) cassettes. For batch culture evolution experiments, standard rich media (10 g*/*L Yeast Extract, 20 g*/*L Peptone and 20 g*/*L Dextrose) was used and was prepared by dissolving 50 g*/*L from a premixed batch of ingredients (Sigma-Aldrich) in H_2_O. For chemostat evolution experiments the same premix was used, but supplemented with 19g/L extra Yeast Extract and 9.5 g/L extra Peptone to obtain a final dextrose concentration of 1 g*/*L. 0.1 mg*/*mL Ampicillin was added to the chemostat media as a safeguard against bacterial contamination. Microscopy experiments were performed in Synthetic Complete (SC) media prepared from Complete Supplement Mixture without amino acids, riboflavin and folic acid (750 mg/L), Yeast Nitrogen Base (6.9 g\L) and either Dextrose (2%w/v) or Ethanol (3%v/v) as a carbon source. All media was filter sterilized to avoid degradation of components during autoclaving.

### Experimental Evolution of Continuous Cultures

#### Multiplexed Chemostat Array Set-Up

We performed our evolution experiments in a dextrose limited chemostat environment by setting up a multiplexed chemostat array of 16 cultures according to the protocol from Miller, Befort (65). YP 0.1%D media was filter sterilized directly into a 10 L glass carboy. During the run, fresh media was provided to the cultures from this carboy by using a peristaltic pump fitted with Marprene tubing. The correlation between rotation speed and media flow rate was empirically determined by measuring the effluent volume at different rotation speeds. Aquarium pumps were used to maintain the positive pressure inside the culture chambers required for the removal of excess culture volume, to keep the cultures aerated and mixed. To minimize evaporation and maintain sterility, air from the pumps was first routed through gas washing bottles and 0.45 µm PFTE filters before entering the culture chambers. The temperature was regulated at 30 °C using heat bocks.

#### Initialization of Multiplexed Arrays

We initialized our multiplexed chemostat arrays by allowing the culture chambers to fill with media until the volume exceeded 20 mL. We dissolved cells from a glycerol stock in YP 0.1 %D media and used to inoculate the cultures by aseptically injecting 4 mL into each culture chamber. In total, 14 *bem3*Δ*nrp1*Δ cultures and 2 WT cultures were inoculated using this procedure. With the peristaltic pump turned off and the aquarium pumps turned on, the *bem3*Δ*nrp1*Δ cultures were left to grow for 4 days and the WT cultures were left to grow for 2 days until they reached saturation (batch phase growth). After the cultures reached saturation, the culture volume was set at 20±1 mL while performing the zero time point sampling.

#### Sampling Regimen

All cultures were sampled twice a week. Samples were taken by replacing the effluent bottles with sterile sampling bottles and collecting the effluent on ice over a period of approximately 2 hours. Directly after sampling, 1 mL of each collected sample was mixed with 500 µL glycerol and stored at *-*80°C. Optical Density (OD) measurements at 600 nm were taken of each sample in 10 mm plastic cuvettes using a photospectrometer (Nanorop 2000C). When necessary, samples were diluted with YP to obtain a final OD of between 0.1-1.5. All samples were diluted in the same media used for blanking the photospectrometer. Effluent volumes were measured daily with a graduated cylinder from which the volume could be read with 0.5 ml precision. On days that sampling took place, the effluent volume of samples was determined after the standard procedure for sampling (glycerol stocks and OD measurements).

#### Calculation of Dilution Rates and Generation Times

We calculated the dilution rate *D* of each sample in our multiplexed chemostat array from the effluent volume using the following formula:

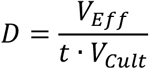

Here, *V*_*Eff*_ is the measured effluent volume, *t* is the time that has passed since the last sampling and *V*_*Cult*_ is the culture volume. At steady state, the growth rate equals the dilution rate [63], allowing the number of generations *G* that have passed to be calculated by:

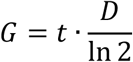

### Experimental Evolution of Batch Cultures

Batch culture evolution experiments were started with 10 *bem3*Δ*nrp1*Δ and 2 WT cultures. The cultures were derived from a single *bem3*Δ*nrp1*Δ and a single WT liquid culture initiated from a glycerol stock and grown to saturation for 2 days in YP 2 %D in a roller drum (set at 40 RPM) at 30°C. After the cultures reached saturation, 10 µL of each starter culture was diluted into 10 mL of fresh YP 2 %D and were placed back into the roller drum. The cultures were diluted by 10 µL into 10 mL of fresh YP 2 %D every 24±2 h. After each dilution, the OD at 600 nm of the remaining culture was measured using the same procedure as described above for the chemostat evolution experiment. Because batch cultures involve frequent population bottlenecks that can reduce genetic variation and possibly purge beneficial mutations (31, 66), it might take longer for an adaptive mutation to settle in the population. To compensate for this effect, the number of generations that the populations were evolved in a batch culture setting was increased to 300 generations (an additional 230 generations compared to the populations evolved in a continuous culture).

### Growth Curve Measurements

Growth curves were obtained by measurements using a plate reader (Tecan Infinite 200 Pro). Cells were inoculated from a glycerol stock in YP 0.1 %D liquid media and grown to saturation for 2 days in a roller drum at 30 °C. On the day of the measurement, the saturated cultures were diluted 1000X into either fresh YP 0.1 %D or fresh YP 2 %D, depending on whether we wanted to measure the fermentation doubling time *T*_*F*_ or the respiration doubling time *T*_*R*_. 100 µL of this culture was pipetted into each well of a sterile 96-well plate (Nunc™Edge 2.0, Thermo Scientific™) with the edge moats filled with 1.7 mL of sterile H_2_O. Each plate contained multiple technical replicates of each sample. As a control for contamination and to allow for background subtraction for downstream processing, 8 wells were filled with blank medium. Measurements were taken during incubation at 30 °C in the plate reader using the following protocol: First, the cells were shaken for 1000 s (linear shaking, 1 mm amplitude) without measurement. After this, the absorbance of each well was measured every 7 min with intermittent shaking (260 s, linear, 1 mm amplitude) for 48 h.

### Growth Parameter Calculations

Doubling times for growth in a glucose environment *T*_*Glu*_ and an ethanol environment *T*_*Eth*_ were extracted from the growth curve measurement in YP 2 %D and YP 0.1 %D, respectively. First, the measured OD values were blanked using the time average value of one of the wells containing blank media. Then, the data was converted to semi-log data by taking the natural logarithm of the blanked OD values. A home written MATLAB script was used to fit a line to the linear portion of the semi-log data to obtain the growth rate in either a glucose or an ethanol environment (Figure S1). These growth rates were converted into doubling times using the following relation:

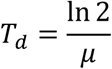

Where *T*_*d*_ is the doubling time corresponding to the growth rate *μ* obtained from the slope of the linear fit. The standard deviation in the distribution of values obtained for *T*_*d*_ was calculated using the MATLAB function std, which uses the following equation:

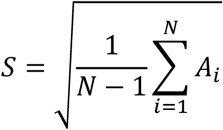

Where *S* is the standard deviation in the observations, *N* is the number of observations and *A*_*i*_ is the *i*_*th*_ observation for *T*_*d*_. The Standard Error of the Mean (SEM) was obtained by dividing *S* by the square root of the total number of observations *N*.

### Microscopy & Microfluidics

Cells were grown to log phase in SC media containing 2% dextrose. Clumps of cells were dissociated prior to imaging by sonicating (Q500 Sonicator, QSonica) in a sealed Eppendorf tube using a cup horn at 70% amplitude for 2 minutes (cycle of 30 seconds pulse on, 15 seconds pulse off). After sonication, each sample was diluted to the same optical density in fresh Synthetic Complete media containing 2% dextrose. Cells were trapped in a microfluidic culture chamber (CellASIC ONIX Y04C-02, Merck – Millipore) after flushing the culture chambers with fresh media for 20 minutes using a pressure of 8 psi. Brightfield images were taken with a Nikon Eclipse Ti-E inverted Microscope using a 60x objective (Plan Apo λ 60X oil, NA: 1.40) with 1 minute intervals. During imaging, cells were maintained in a constant flow of media using a pressure of 1 psi. Cells were subjected to a media switch by changing from an inlet with SC media containing 2% dextrose to an inlet with SC media with 3% ethanol after 8 hours of imaging.

### DNA extraction, Illumina library preparation & Whole Genome Sequencing

We extracted genomic DNA from liquid cultures grown for two overnights for each of the 16 chemostat samples, 10 serial dilution samples and a non-evolved yLL132a ancestor with the MasterPure Yeast DNA Purification Kit (Epicentre, Madison, WI, USA) following the manufacturer’s protocol. We included a RNase A (Qiagen, Hilden, Germany) treatment step in the protocol and collected DNA in a final volume up to 30 µL H_2_O. We pooled up to three extractions per sample using the Genomic DNA clean & Concentrator kit (Zymo Research, Irvine, CA, USA), following the supplied protocol. We eluted DNA in a final volume of 30 µL. We assessed DNA quality by 0.8 % agarose gel electrophoresis and quantity by fluorometry using a Qubit 4.0 Fluorometer (Invitrogen, Carlsbad, CA, USA). Samples were individually barcoded and pooled into a single library with the NEB Next Ultra DNA Library Prep Kit (New England Biolabs, Ipswich, MA, USA) and sequenced on a HiSeq machine (Illumina, San Diego, CA, USA) by Novogene (Bejing, China).

### WGS Data analysis

We first checked raw paired-end reads (150 bp) for quality with the FASTQC toolkit (version 0.11.7; http://www.bioinformatics.bbsrc.ac.uk/projects/fastqc). We removed low quality ends (Quality scores < 20; and first 9 bases of all reads), and re- moved duplicates with the FastX toolkit (version 0.0.14; http://hannonlab.cshl.edu/fastx_toolkit/). We downloaded the R64-1-1 *S. cerervisiae* genome from the Saccharomyces Genome Database (SGD; www.yeastgenome.org; September 2018) and used it as our reference. We indexed the reference genome with the Burrows-Wheeler Aligner (BWA; version 0.7.17; (Li and Durbin, 2010)), and SAMtools (version 1.8; (Li, 2011; Li and Durbin, 2009)), and generated a dictionary with Picard (version 2.18.5; https://broadinstitute.github.io/picard/). We mapped sequences from all samples individually to the reference with BWA-MEM sorted and indexed mapped reads into a BAM file with SAMtools. We performed multisample SNP calling and additional indexing with SAMtools and BCFtools (version 1.8). We plotted and checked statistics, e.g., TS/TV and quality of sites and read depth, with BCFtools. These statistics were used to filter out SNPs and Indels with low quality sites (QUAL > 30), low read depth (DP > 20), and variants in close proximity to gaps (SnpGAP 10). We annotated the VCF file with snpEff (version 4.3T; (Cingolani et al., 2012)) with R64- 1-1.86. We then retrieved variants (SNPs and indels) of interest through comparison of variants between the reference strain, the ancestor strains, and the evolved strains. We excluded variants that were only different between R64 and all our W303 samples, as these merely display differences between the two types of strains (see e.g., (Ralser et al., 2012)). Synonymous variants, variants in noncoding regions, and stop retained variants were excluded. Mutations in telomeric regions and in Long Terminal Repeats (LTRs) were excluded from analysis due to the natural variation that occurs in the genomic sequence of these regions. To find causative mutations, we looked for genes that mutated in at least two evolved lines, excluding those that appeared only in the mutant line(s) from one environment and the wild-type line(s) of the other environment. From the resulting list of genes, genes corresponding to dubious or uncharacterized Open Reading Frames (ORFs) were removed according to their description on SGD. Two genes (*RPS29B* and *ECM33*) had acquired the same mutation across all 22 parallel evolved *bem3*Δ*nrp1*Δ lines that sweeped the population, suggesting that these mutations were acquired in the ancestor before the different cell lines were split. Although these mutations might have some fitness benefit in the *bem3*Δ*nrp1*Δ background, they do not explain the adaptation we observe during our evolution experiments and we therefore excluded them from further analysis.

## Supplementary Material

**Figure S1:**
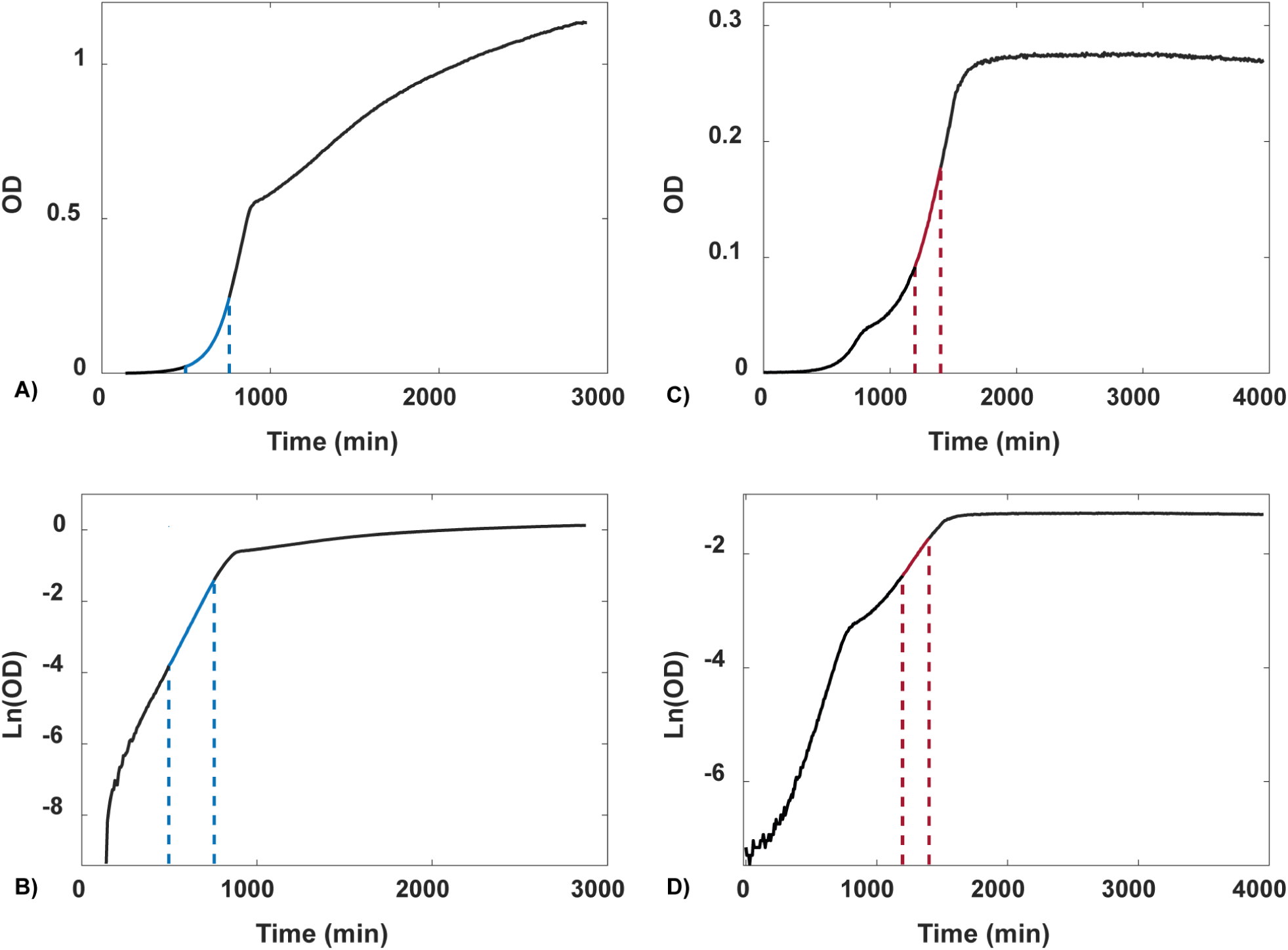
Low glucose media induces earlier diauxic shift, allowing better quantification of growth in respiration. (**A-B**) When grown on high glucose (2%) media, most of the biomass is produced by fermentation. (**C-D**) Lowering the glucose concentration to 0.1% induces an earlier diauxic shift and improves visualization and quantification of growth in respiration. Dashed lines indicate the period of exponential growth for growth in YP+2% glucose (blue) and for growth in YP+0.1% glucose (red).

Although the diauxic shift phase of growth is clearly visible in both cases as a temporary cessation of growth (red dots, Figure 1A,B), the high OD at which the diauxic shift occurs in 2% glucose media makes it unsuitable to quantify growth beyond this point due to the possible non-linear relationship between cell density and absorbance at high OD values. Conversely, in 0.1% glucose media the diauxic shift occurs at a relatively low density, decreasing the amount of signal relative to the noise during pre-diauxic growth. Therefore, growth rates in terms of population doubling times in a glucose environment (*T*_*Glu*_) were determined from growth data in 2% glucose media, while growth rates in an ethanol environment (*T*_*Eth*_) were obtained from growth data 2% glucose media.

**Figure S2:**
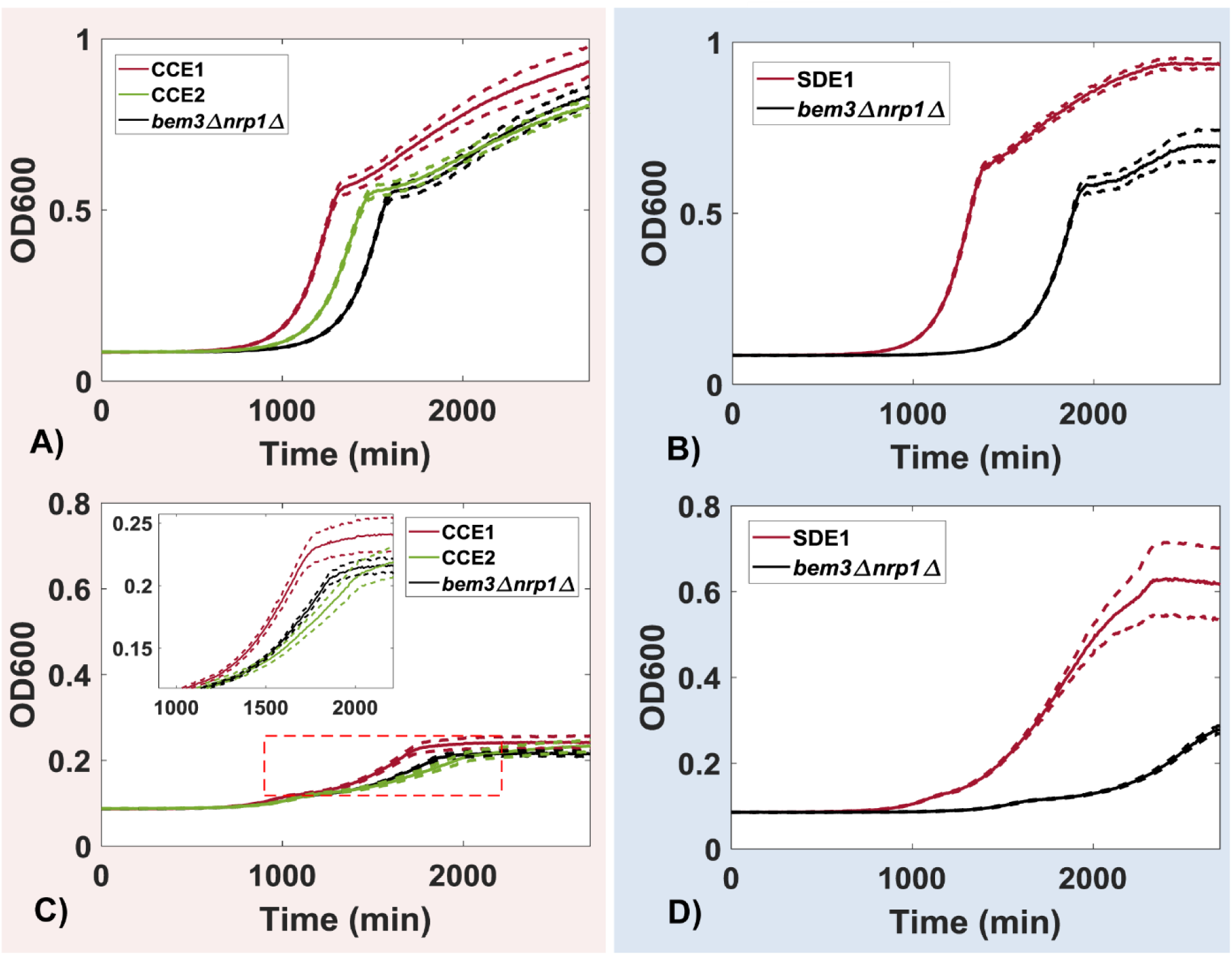
Growth curves of selected evolved lines CCE1, CCE2 and SDE1. (**A**,**C**) Bulk measurements in (**A**) 2% dextrose and (**C**) 0.1% dextrose media of selected evolved mutant lines CCE1 and CCE2 from the continuous culture experiment together with the mutant ancestor strain. (**B**,**D**) Bulk measurements in (**B**) 2% dextrose and (**D**) 0.1% dextrose media selected evolved mutant line SDE1 from the batch culture experiment. Dashed lines display the SEM.

**Figure S3:**
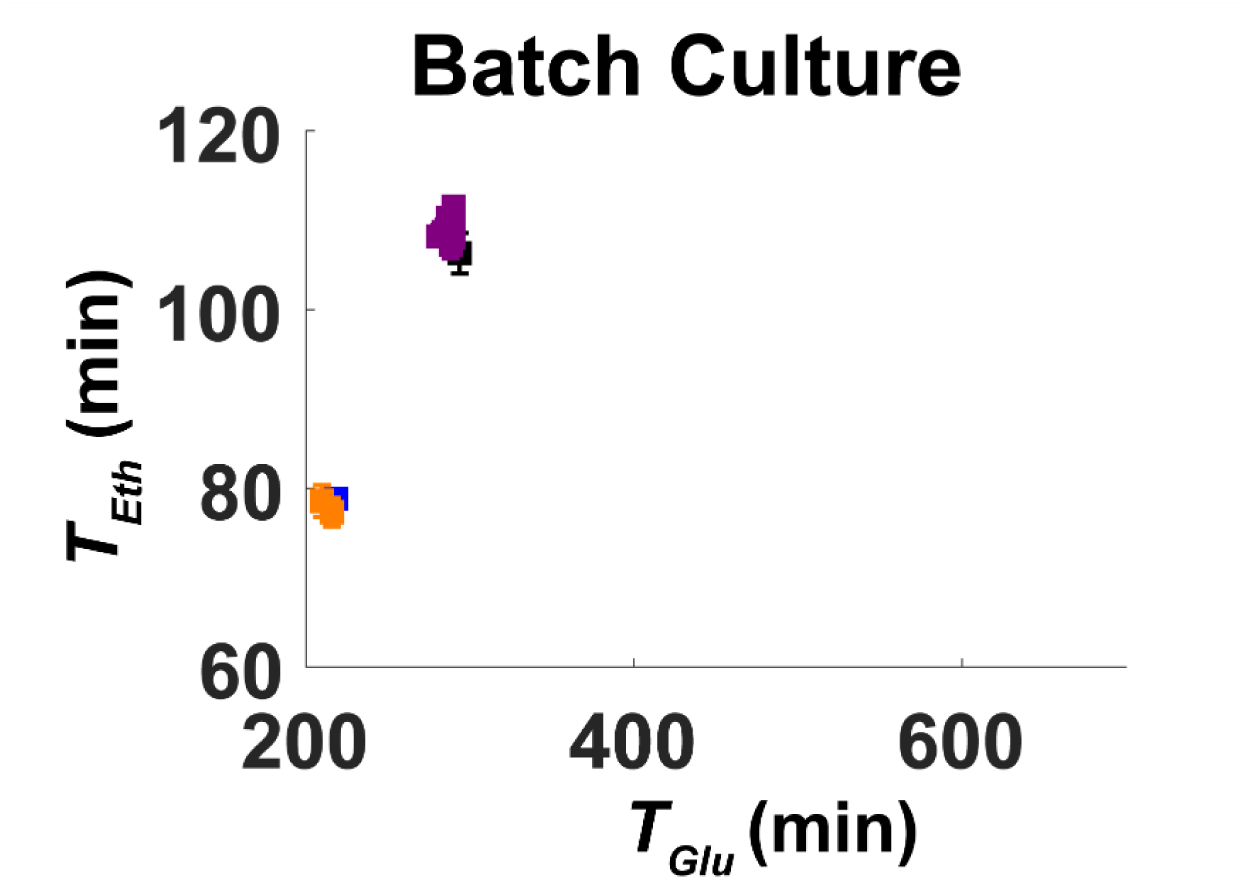
Scatter plot of *T*_*F*_ vs *T*_*R*_ after 70 generations in a batch culture. After 70 generations of evolution in a batch culture the evolved cell lines were still phenotypically highly similar to their ancestral strains. This was the case for both the *bem3Δnrp1Δ* mutants and the WT strains.

**Figure S4:**
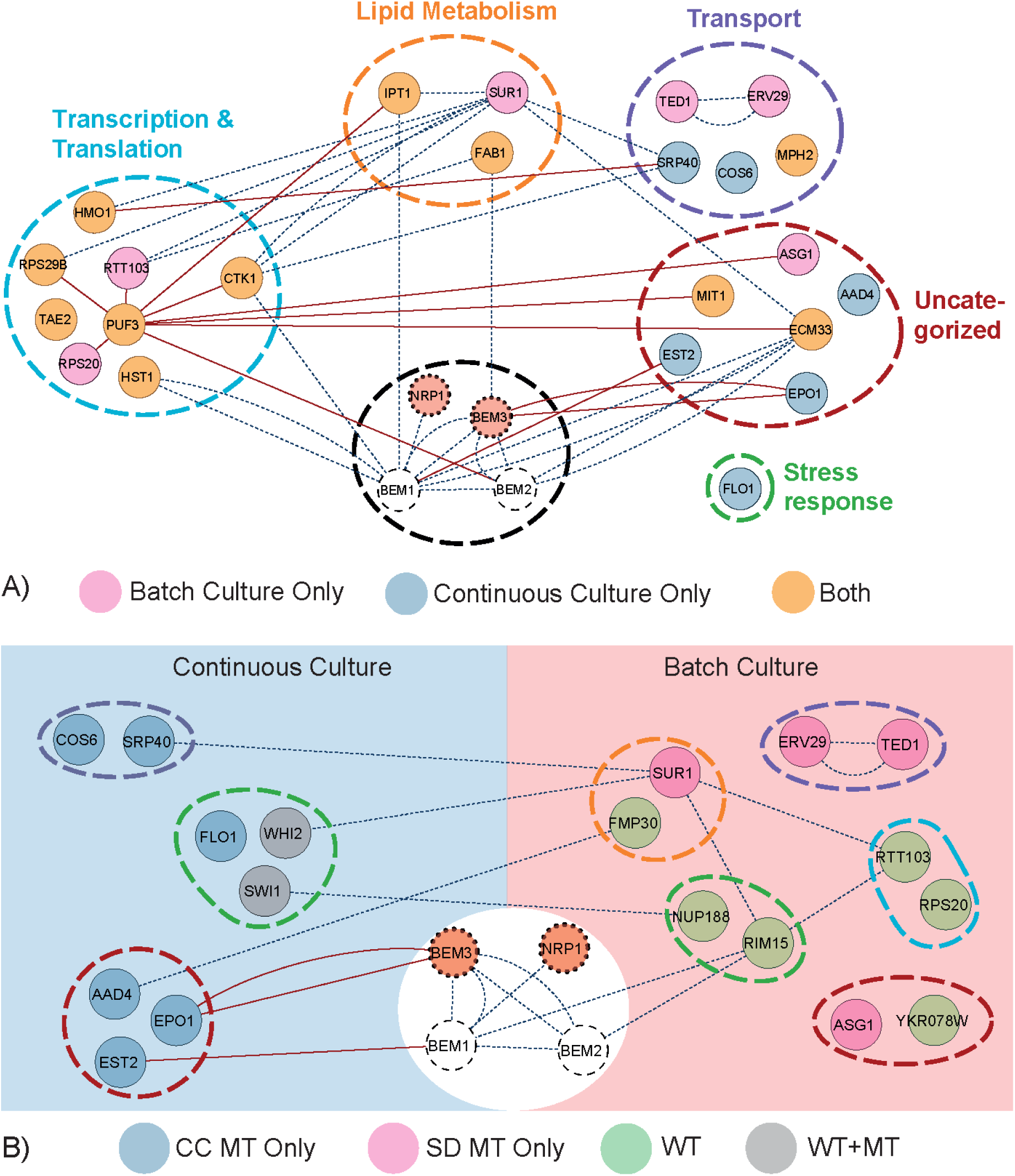
Mapping of the genes mutated in during the evolution of *bem3Δnrp1Δ* mutants to their cellular process GO-terms. (**A**) Genes mutated in *bem3Δnrp1Δ* mutants, but not WT, after evolution are grouped according to their cellular process GO-term. Solid red lines display physical interactions and dashed blue lines display genetic interactions. The interaction network affected by a *bem3Δnrp1Δ* mutation that was described by [8] is included (black dashed circle). (**B**) Genes mutated in the evolved lines, including the WT lines, that are specific to a continuous culture or a batch culture environment. Color coding is the same as in (**A**).

### Changes in respiration and fermentation rates do not result from a trade-off with the rate of metabolic switching

Why do some of the *bem3Δnrp1Δ* lines evolved in a continuous culture compromise on *T*_*Eth*_? One possibility is that this is due to pleiotropic effects, where some of the mutations that cause improvements in *T*_*Glu*_ are detrimental for *T*_*Eth*_. Such mutations are able to sweep through the population in the constant environment of a continuous culture due to the lack of selection for *T*_*Eth*_. However, an alternative possibility is that there exists a trade-off between *T*_*Eth*_ and another parameter that is selected for without our knowledge. We decided to investigate the possibility of a trade-off for the diauxic lag-phase, as various studies have found a trade-off between the maximal growth rate of cells and the rate of switching between nutrients during diauxic growth (67, 68).

Although the cells in a continuous culture are predominantly in a fermentative state they might be close to exhausting the glucose source (Table 1) (31, 37). Despite that it has been demonstrated by models to be unlikely to occur during growth in continuous culture (69), we did not want to exclude the possibility that some populations might have adopted a strategy where increasing the switch rate switch between fermentation and respiration is beneficial. We therefore explored the option of a trade-off for the diauxic switch rate *T*_*S*_ (Figure 3A). The value of *T*_*S*_ was determined geometrically: The linear fits for *T*_*Glu*_ and *T*_*Eth*_ were taken and the horizontal line that connects these two fits while crossing the largest number of datapoints was determined (Figure 4B). We found that this procedure typically leads to a good measure for *T*_*S*_, although the error in the estimate increased for datasets with very short diauxic lag times (Figure S6), which results in a bias for larger values of *T*_*S*_.

We plotted the obtained values of *T*_*S*_ against the values of *T*_*Glu*_ (Figure 3C,D) and *T*_*Eth*_ (Figure 3E,F). We first looked if we could identify a trade-off between these different growth parameters for our populations evolved in a continuous culture (Figure 3C,E). In the case of a trade-off we would expect a higher value for either *T*_*Glu*_ or *T*_*Eth*_ (corresponding to slower growth in fermentation and respiration, respectively) to be accompanied by lower values for *T*_*S*_ (corresponding to a faster diauxic switch. However, the results show no clear correlation for the populations evolved in a continuous culture, neither between *T*_*S*_ and *T*_*Glu*_ nor *T*_*Eth*_ (Figure 3C,E). Instead there appears to be no restriction for the evolution of higher or lower values for *T*_*S*_. Performing the same analysis for populations evolved by serial dilutions show that this environment also does not display any trade-off between *T*_*S*_ and *T*_*Glu*_ or *T*_*Eth*_ (Figure 3D,F). Instead, there appears to be selection for adaptations that improve all three parameters *T*_*S*_ and *T*_*Glu*_ or *T*_*Eth*_ compared to their ancestor, consistent with the observation that all three traits are under selection in this environment.

These results show that the decrease in *T*_*Glu*_ observed in the lines evolved in a continuous culture (Figure 2A) does not result from a trade-off with *T*_*S*_, but is likely to be a consequence of pleiotropy: some mutations that improve *T*_*Glu*_ affect the network that regulates *T*_*Eth*_, although this effect can be both positive and negative. Selecting on both parameters allows only those mutations that improve both *T*_*Glu*_ and *T*_*Eth*_ to settle in the population. Furthermore, this demonstrates that the adaptations that result in the observed changes in *T*_*Glu*_ are pleiotropically linked to networks that affect *T*_*Eth*_, but not to networks that affect *T*_*S*_.

**Figure S5.**
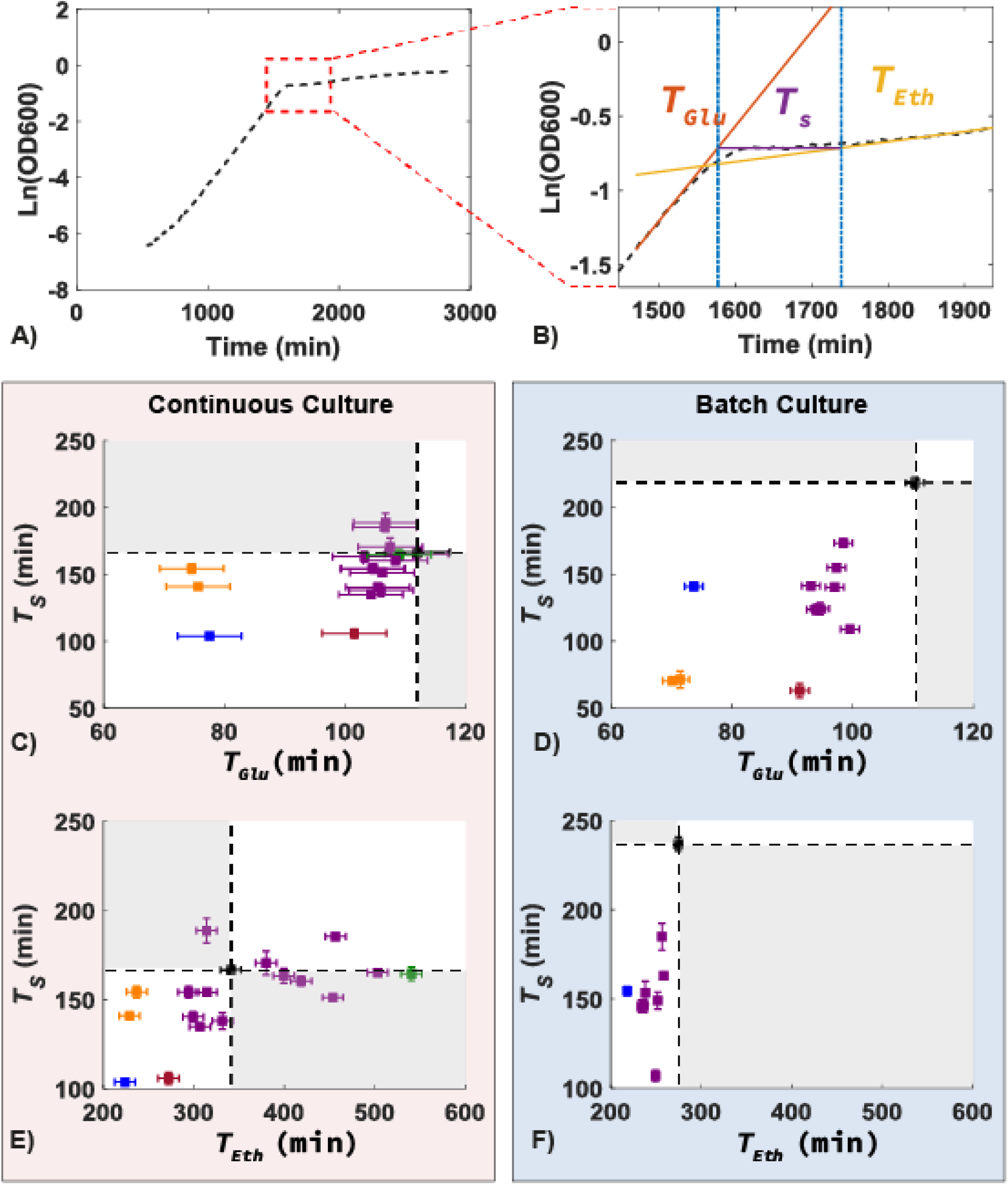
Evolution in both environments is not restricted due to trade-offs with the diauxic lag phase. (**A**) Semi-log plot of an example growth curve used to estimate *T*_*S*_. A zoom-in of the area annotated by the red box is shown in (**B**). (**B**) Schematic showing the procedure to determine the length of the diauxic lag phase. The time to switch between fermentative growth (*T*_*Glu*_) and respiratory growth (*T*_*Eth*_) was obtained by finding the two intersection points of the best fitted horizontal line during the diauxic shift (purple) and the lines used to determine the doubling time *T*_*Glu*_ (red) and *T*_*Eth*_ (yellow). (**C-D**) Scatter plot showing *T*_*S*_ vs *T*_*Glu*_ for the ancestral and lines evolved in (**C**) a continuous culture and (**D**) lines evolved by serial dilution. The plot shows no correlation between *T*_*S*_ and *T*_*Glu*_. (**E-F**) Scatter plot showing *T*_*S*_ vs *T*_*Eth*_ for the ancestral and lines evolved in (**E**) a continuous culture and (**F**) lines evolved by serial dilution.

**Figure S6:**
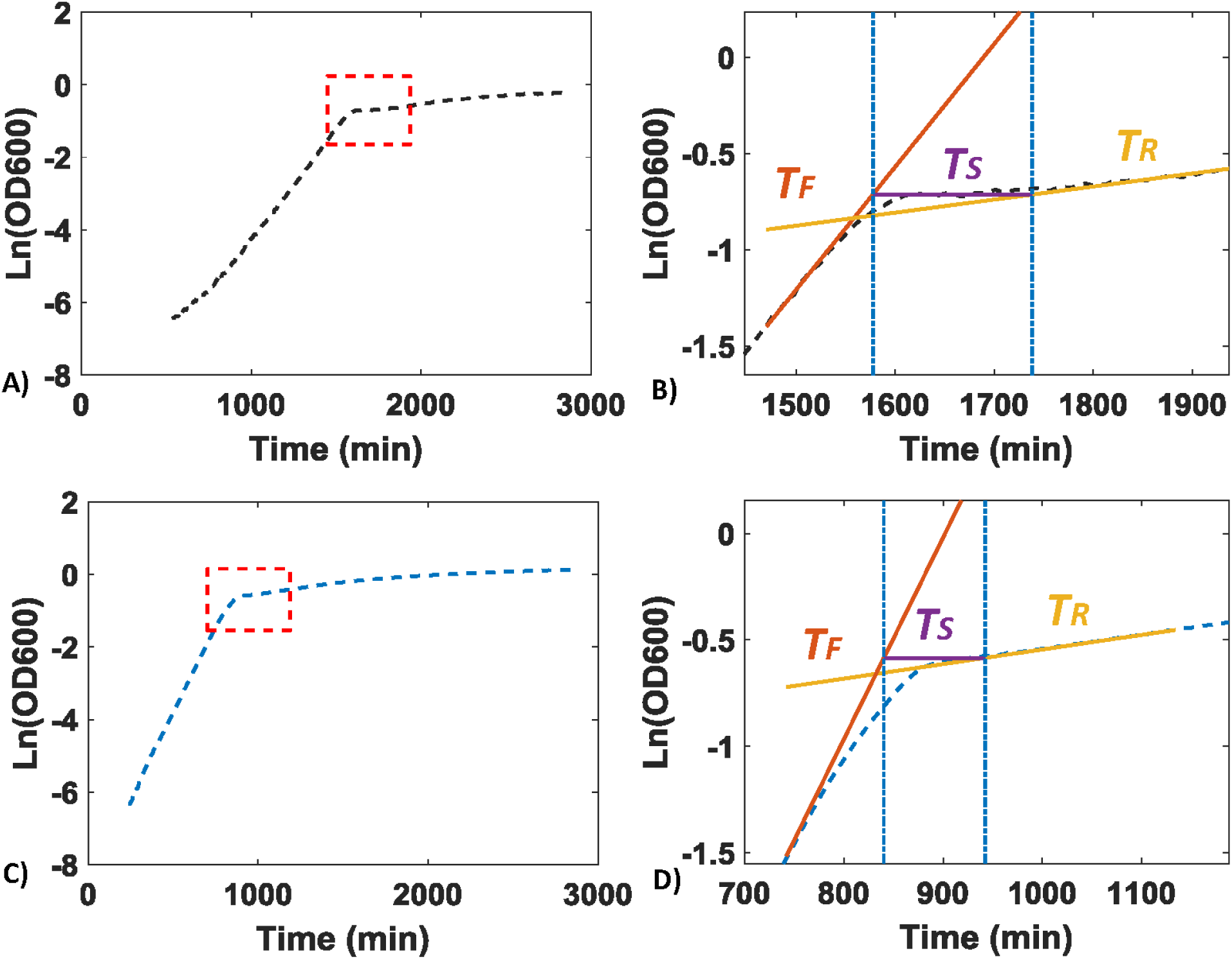
The error in the estimate for *T*_*S*_ increases for shorter values of *T*_*S*_. Figure shows the increase in the error when going from long switching times (**A-B**) to shorter switching times (**C-D**).

**Table S1:** Glucose and ethanol concentrations (mM) in the residual media during growth of a WT strain and a *bem3Δnrp1Δ* mutant in log phase, a continuous culture and stationary phase. BDL: Below Detection Limit

| Glucose (mM) | Log Phase | Continuous Culture | Stationary Phase |
| --- | --- | --- | --- |
| WT | 6.079 | 0.023 | BDL |
| *bem3Δnrp1Δ* | 6.737 | 0.036 | BDL |
| Ethanol (mM) | Log Phase | Continuous Culture | Stationary Phase |
| WT | 2.294 | 1.373 | BDL |
| *bem3Δnrp1Δ* | 1.301 | 0.66 | 0.786 |

Comparison of the residual glucose and ethanol concentrations of our continuous culture to those of early log phase and late stationary phase cultures in glucose limited media (0.1% w/v) using HPLC. This confirmed the presence of low concentrations of both glucose and ethanol in the continuous culture. Specifically, glucose is present at a concentration lower than in early log phase, but not yet fully depleted as is the case in late stationary phase. The ethanol concentration shows the same pattern, but only for the WT strain: In a stationary phase culture of *bem3Δnrp1Δ* cells, there was still a detectable amount of ethanol present. This is consistent with the previous observation that these cells are inefficient in passing through the diauxic shift (Figure 1E), making them unable to fully consume the ethanol through respiration. This portrays the advantage of being phenotypically plastic in a batch culture, as it allows continued growth on an otherwise unused carbon source. In contrast, a continuous culture supports constant growth without requiring passage through the diauxic shift (69).

## Notes

### Competing Interest Statement

The authors have declared no competing interest.

